# Effects of human stress on horse behavior and interaction quality

**DOI:** 10.64898/2025.12.03.692157

**Authors:** Yeonju Choi, Youngwook Jung, Carissa L. Wickens, Minjung Yoon

## Abstract

Human-horse interaction can be influenced by the emotional and physiological state of humans, as horses are highly sensitive to human stress. Understanding these effects is essential for improving outcomes in equine-assisted activities. This study examined how human stress impacts horse behavior and interaction quality. Seven humans and seven horses participated in a crossover design experiment in which participants viewed either stressful or relaxed video stimuli prior to interacting with horses. Electroencephalography (EEG), heart rate variability (HRV), and salivary cortisol was collected to assess human physiological responses, while the Human-Animal Interaction Scale (HAIS) and an ethogram evaluated interaction quality and horse behaviors. Stressful stimuli significantly increased beta and high-beta activity and altered HRV indices, indicated by increased low-frequency power and decreased high-frequency power. HAIS scores were lower under stress conditions, reflecting reduced interaction quality. Horses exhibited more pushing behavior and less lip-licking when handlers were stressed, although task completion time did not differ between conditions. These results demonstrated that human stress negatively influences both physiological states and behavioral outcomes within human-horse interactions. Therefore, managing emotional states and stress levels may enhance interaction quality and effectiveness in equine-assisted programs.

## 1. Introduction

Human-horse interactions have been a subject of interest due to their potential therapeutic and emotional benefits. The bond between humans and horses has been explored in various contexts, including therapy [1], sports [2], and recreational activities [3]. Recent research has focused on understanding how human emotions and stress levels can influence horse behaviors, as horses are known for their sensitivity to human cues and emotions [4, 5]. This sensitivity is thought to play a crucial role in the human-horse relationship, affecting both the emotional state of the horse and the outcomes of interactions.

Stress is a complex physiological and psychological response to perceived threats or challenges, and it can significantly affect human behavior and cognition [6]. In human-animal interactions, stress level of humans can be communicated to the animal, affecting the behavior and emotional state of animals [7, 8]. This is particularly relevant in human-horse interactions, as horses are highly perceptive and responsive to subtle changes in human body language and emotional cues [9, 10].

Electroencephalography (EEG) is a non-invasive method used to record electrical activity of the brain [11]. By placing electrodes on the scalp, EEG captures brainwave patterns that reflect states of arousal, attention, and relaxation. In this study, specific EEG frequency bands, such as alpha, beta, high-beta, and theta waves are analyzed to evaluate stress levels. Alpha waves are typically associated with states of relaxation, calmness, and a wakeful rest [12]. Beta waves are associated with active thinking, focus, and alertness. An increase in beta wave activity is often observed during states of stress, anxiety, or high mental arousal [13]. High-beta waves are linked to intense concentration, high cognitive load, and arousal. They are typically seen during periods of stress and anxiety [14]. Theta waves are related to deep relaxation and the early stages of sleep. While theta waves are generally associated with relaxation, their role in stress is more complex. In some cases, increased theta activity can occur during states of heightened emotional processing or when the brain is trying to cope with stress [15].

Heart Rate Variability (HRV) is widely used to measure stress because it reflects the activity of the autonomic nervous system, which controls involuntary bodily functions, including heart rate (HR) [16]. HRV is sensitive to both acute and chronic stress. Acute stress, such as an immediate threat or emotional challenge, leads to a rapid decrease in HRV due to increased sympathetic activity. This sensitivity makes HRV a useful tool for assessing the physiological impact of stressors.

Cortisol is a steroid hormone released by the adrenal glands in response to stress [17, 18]. It is a key biomarker for the activation of the hypothalamic-pituitary-adrenal axis, which is the central stress response system. Measuring cortisol levels can provide insights into the physiological stress response in both humans and animals. In this study, cortisol levels were used to objectively assess stress in both humans and horses, complementing the HRV and EEG data.

This study aimed to assess the influence of human stress levels on horse behaviors by evaluating physiological and behavioral responses in both humans and horses to understand how the stress is perceived and responded to by horses. We hypothesized that human stress, induced by exposure to stressful stimuli, lead to observable changes in horse behavior, reflecting the sensitivity of horses to their handlers’ emotional states.

## 2. Materials and Methods

### 2.1. Experimental subjects

This study involved seven human participants (4 females and 3 males), with an average age of 27.8 years. All participants were healthy adults with no reported mental illness. These experimenters were informed about the experimental procedure, and they also received additional instructions on the experimental day. The equine subjects consisted of seven horses, including 5 Thoroughbreds, 1 pony, and 1 Warmblood, with an average age of 8.9 years. All the horses were in good health and had experience with basic handling and groundwork exercise. Each horse was housed individually in a stall and received four feedings per day, consisting of 1.5% of their body weight in Timothy hay and 0.5% of their body weight in commercial concentrates (Cavalor pianissimo 20 kg, Cavalor, Belgium). They allowed free access to water.

### 2.2. Experimental design

This study was performed at Gyeongbuk Natural Science High School, Sangju, Republic of Korea. The experiment was conducted using a crossover design, where each participant was exposed to two different conditions: a stressful video and a relaxed video. The study was divided into three main phases for each condition: baseline, after watching videos, and after horse-related activity. Baseline data was collected before video exposure and any activities with the horses, serving as the control measurement. Then measurements were taken after the participants watched the assigned video but before interacting with the horse. Finally, measurements were collected after the participant completed a series of activities with the horse. Each participant was assigned to one horse and went through the experimental process over the course of a single day, with one session in the morning and another in the afternoon. The activities with horses were conducted in 40 m Ⅹ 60 m indoor arena included leading, backing up, walking side-to-side, crossing over tarp obstacles, touching a gym ball, following the previous study [19].

### 2.3. Behavior observations and assessments

The horse behaviors were recorded during activities using action camera (AC10, Thinkware, Republic of Korea). To evaluate, an ethogram with different behaviors, including ear pinned back, tail swishing, push, bucking, resisting, body shaking, lip-liking, and blowing (Table 1). This ethogram was constructed based on previous studies [20–23]. The time taken to cross over obstacle course consisting of blue tarps and to touch a gym ball was also recorded during activities. Additionally during the activities, the scores of interactions between handlers and horses were measured via Human-Animal Interaction Scale (HAIS), which provides an assessment device for measuring human-animal interactions [24].

**Table 1.** Ethogram of horse behaviors during interactions with handler.

| Behaviors | Description |
| --- | --- |
| Ear pinned back | Flattening or positioning the ears behind vertical or flat against the head |
| Tail swishing | Lateral, dorsoventral or circular motion of the tail |
| Push | Applying pressure with the neck, shoulder, chest, body to displace the handler |
| Bucking | Sudden arching of the back and leaping, and kicking with hind legs |
| Resisting | Ignoring or resisting the cues provided by the rider |
| Body shaking | Shaking its entire body, starting from the head and moving down to the tail |
| Lip-licking | Moving its tongue to lick the upper or lower lips |
| Blowing | Short non-vocal sound given during exhalation |

### 2.4. EEG

EEG data were collected using QEEG-8FX system (Laxtha Inc., Daejeon, Republic of Korea). EEG signals were processed and analyzed using Telescan software (CD-TS-2.2; Laxtha Inc.). EEG electrodes were placed at the prefrontal (Fp1, Fp2), frontal (F3, F4), temporal (T3, T4), and parietal (P3, P4) sites based on the international 10-20 system. EEG was recorded for 2 min at each timepoint. The raw data underwent band-pass fast fourier transform-filtering (0.5-50 Hz). Absolute power was calculated for the following frequency bands [25]: alpha (8-12 Hz), beta (13- 30 Hz), high-beta (23-40 Hz), theta (4-7.5 Hz). To visualize overall cortical activation, power spectral density values from all electrodes were averaged to produce a global spectral profile.

### 2.5. HRV

HRV was assessed using ECG recordings from the QEEG-8FX system (Laxtha Inc.). Two electrodes were placed on the left and right inner wrists of the participants, respectively. HRV was recorded for 2 min at each timepoint. ECG data were analyzed using Kubios HRV software (version 4.1.1; Kubios Oy, Kuopio, Finland) to measure HRV parameters including HR, low frequency power (LF), high frequency power (HF), and LF/HF ratio.

### 2.6. Cortisol analysis

Saliva samples were collected from both humans and horses to quantify cortisol concentrations. Synthetic saliva collection swabs (Salivette, No.51.1534.500; Sarstedt, Nümbrecht-Rommelsdorf, Germany) were used [26, 27]. Human participants placed the swabs in the mouth for 1-2 min until saturated, while for horses, the swabs were attached to a bit and held in the mouth for 1-2 min. After saturation, swabs were immediately transferred into polypropylene tubes and stored at 4℃ during transportation. Samples were then centrifuged at 1,500 x g for 3 min, and the extracted saliva was stored at −80℃ until analysis. Cortisol concentrations were determined using a commercial ELISA kit (ADI-900-071; Enzo Life Sciences, USA) according to the manufacturer’s protocol. The assay sensitivity was 56.72 pg/mL. The mean intra- and inter-assay coefficients of variation were <10.5% and 13.4%, respectively.

### 2.7. Ethical consideration

The research protocol was reviewed and approved by the Institutional Review Board (permit number 2025-0225) and Animal Experimentation Ethics Committee (permit number 2024-0024, 2024-0043) of Kyungpook National University. Experiments were conducted in accordance with the committee guidelines. Human participants were prospectively recruited between 1 May 2025 and 9 May 2025. All participants were informed about the purpose and procedures of the study prior to participation and provided written informed consent. No minors were included in the study. During the experiments, the researchers carefully handled the horses to reduce the duration of the experiments and prevent any undue stress on the animals.

### 2.8. Statistical analysis

The data collected in this study were analyzed using Prism 10 (Graphpad Software, California, United States). Differences in HRV, brain wave data, and cortisol concentrations in handlers were assessed using a mixed model for repeated measures to compare means between data from different time points during the experiments. Additionally, post-hoc analysis was performed using Tukey’s multiple comparison test. Differences in HAIS scores in handlers and horses, cortisol concentrations in horses, and the frequencies of behaviors shown by horses during activities were analyzed using paired t-tests. P-values less than 0.05 were considered statistically significant.

## 3. Results

### 3.1. Brainwave changes of handlers

The analysis of brainwave activity showed differential responses to stressful and relaxed conditions in terms of beta and high-beta waves, while alpha and theta waves remained relatively unchanged. Alpha wave activity did not show significant changes across the different conditions. The mean values for alpha power in both stressful and relaxed conditions remained relatively consistent throughout the baseline, before, and after activity phases (Fig 1A). Beta wave activity significantly increased after participants watched the stressful video and then decreased following the activities with the horses. In contrast, there were no significant changes observed in beta wave activity after watching the relaxed video (Fig 1B). High-beta wave activity showed a similar pattern to beta waves, with a significant increase after watching the stressful video and a decrease after interacting activities with the horses. No significant changes were observed following the relaxed video (Fig 1C). Theta wave activity remained stable and did not show significant changes across the different conditions. The mean values for theta power consistent before and after watching both stressful and relaxed videos (Fig 1D).

**Figure 1.**
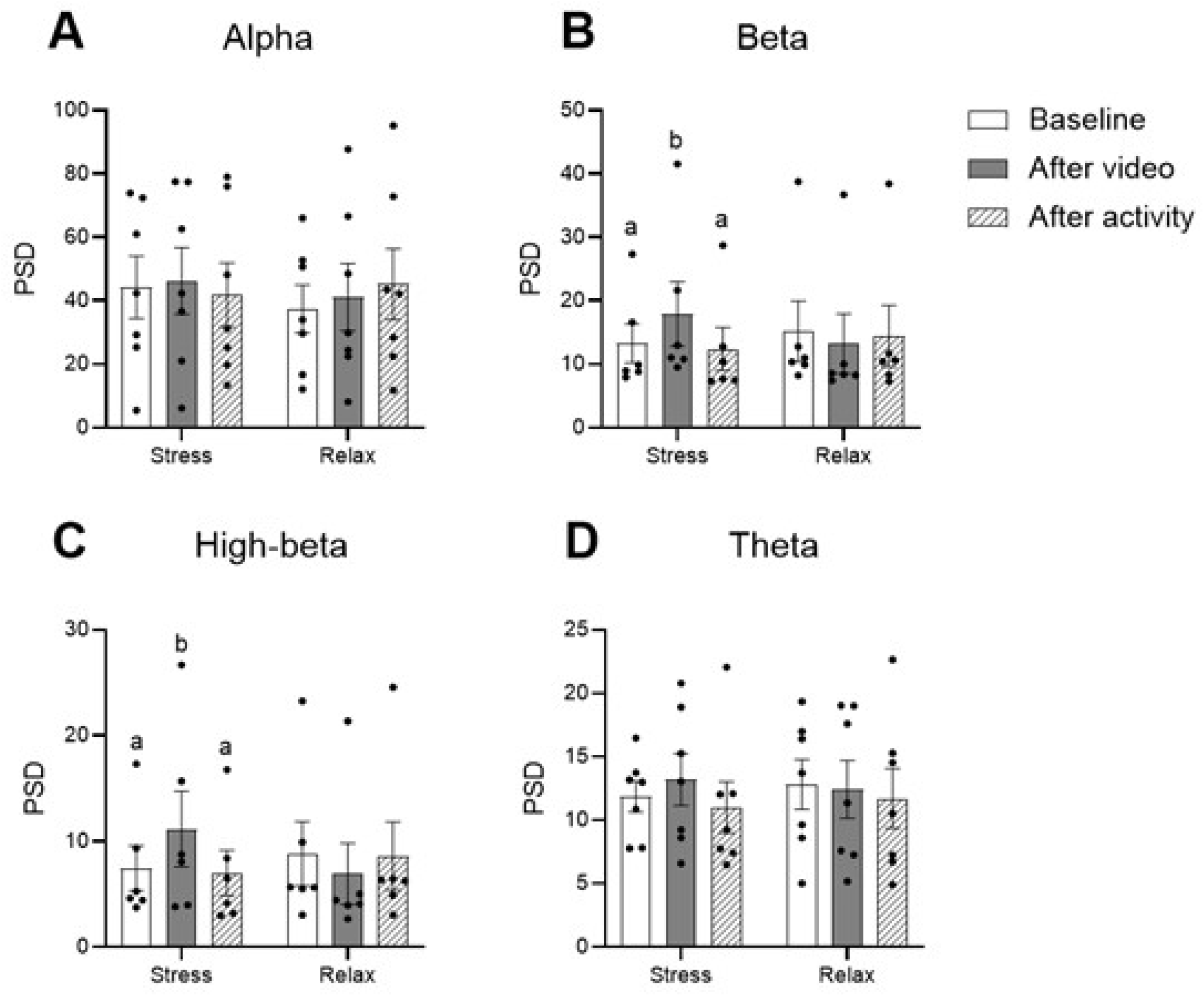
Mean power spectrum density of alpha waves (A), beta waves (B), high-beta waves (C), and theta waves (D) of handlers. Dots indicate individual data points. Alphabets indicate significant differences. PSD, power spectrum density.

### 3.2. HRV changes of handlers

The average HR (beats per minute) did not show significant changes between the different conditions (Fig 2A). LF power significantly increased after participants watched stressful videos, indicating an increase in sympathetic nervous system activity. Following the activities with the horse, LF power decreased (Fig 2B). There were no significant changes in LF power after watching the relaxed video. There was a significant difference in LF powers between the timepoints following the stressful and relaxed videos. HF power significantly decreased after watching the stressful video, indicating a suppression of the relaxation response. After interacting with the horses, HF power increased (Fig 2C). HF power also increased after the relaxed video compared to baseline. Significant differences in HF were found between the stressful and relaxed conditions. The LF/HF ratio showed a similar pattern to LF changes. The ratio significantly increased after watching the stressful video. This ratio decreased after the activities with horses (Fig 2D). No significant changes were noted in the LF/HF ratio after watching relaxed video.

**Figure 2.**
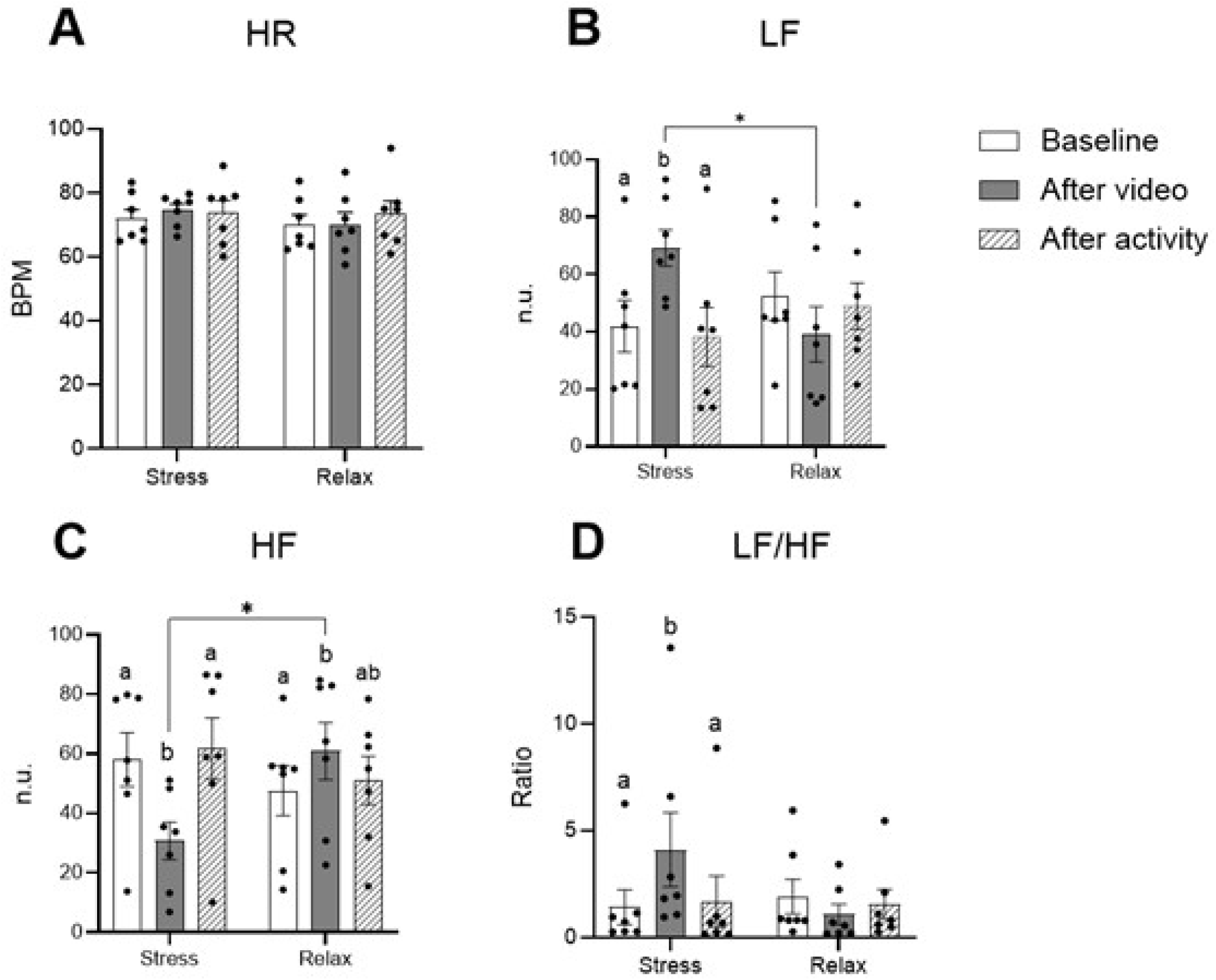
Mean heart rate (A), low frequency band (B), high frequency band (C), and ratio of low frequency and high frequency (D) of handlers. Dots indicate individual data points. Alphabets and asterisks indicate significant differences (* **<** 0.05). HR, heart rate; BPM, beats per min; LF, low frequency; HF, high frequency; n.u., normalized units.

### 3.3. Changes in human-horse interaction and cortisol levels

The HAIS for humans was significantly lower when participants watched the stressful video compared to the relaxed video (Fig 3A). This indicates that the quality of interaction between humans and horses was negatively affected by humans’ elevated stress levels. When participants were in a more relaxed state, as induced by the relaxed video, they tended to interact more positively with the horses. Similarly, the HAIS for horses was significantly lower when their handlers had watched the stressful video compared to the relaxed video (Fig 3B). This pattern suggests that the horses were responsive to the handler’s stress levels, exhibiting less favorable behaviors during interactions when the handlers were stressed.

**Figure 3.**
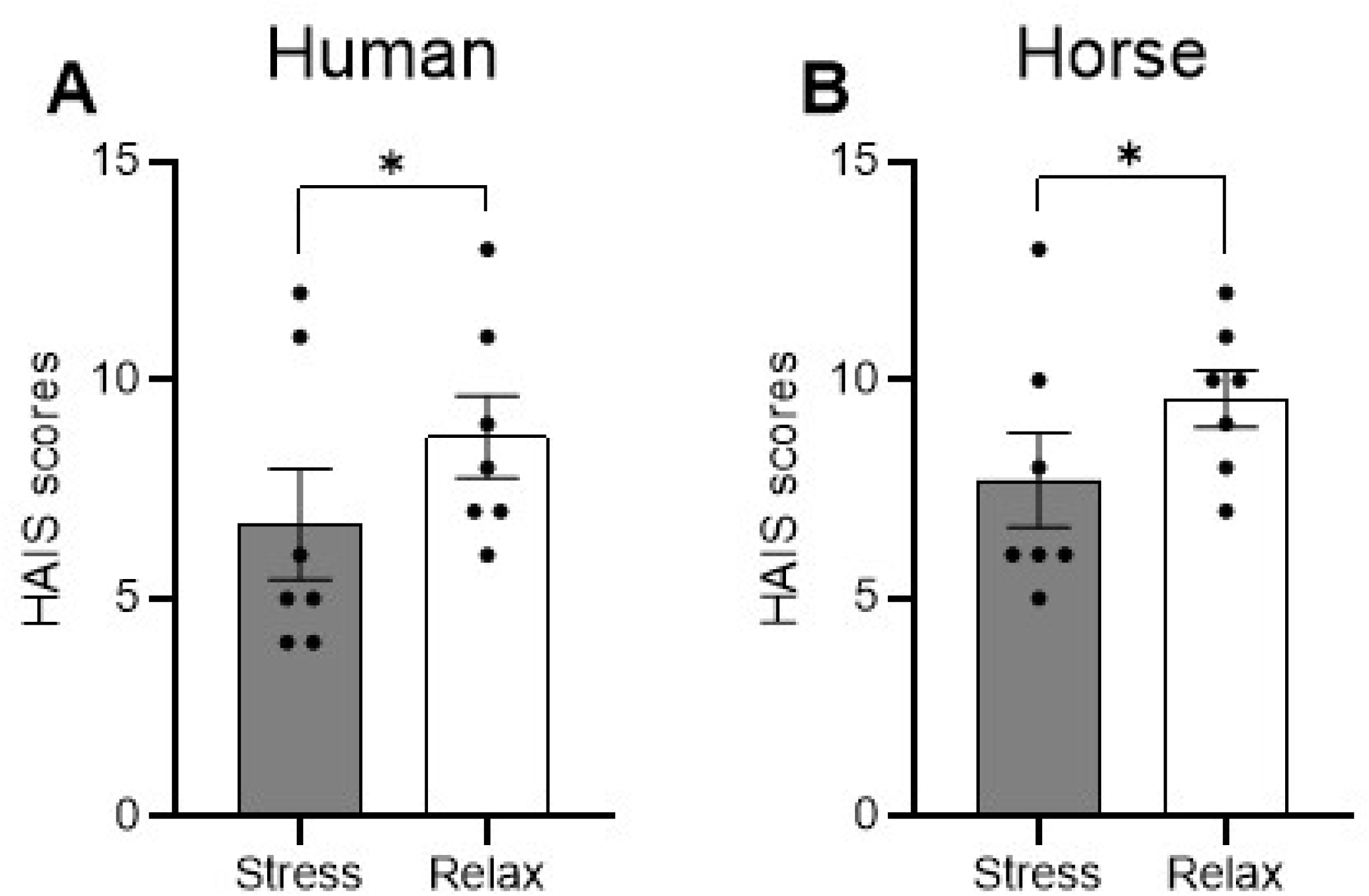
Mean human-animal interaction scores of handlers (A) and horses (B) Dots indicate individual data points. Asterisks indicate significant differences (* < 0.05).

### 3.4. Changes in cortisol concentrations in humans and horses

In humans, cortisol concentrations increased when participants watched stressful video but decreased after participating in horse-mediated activities. In contrast, cortisol levels remained constant when participants watched a relaxing video (Fig 4A). Significant differences were observed between the stressful and relaxed conditions. Unlike humans, horses did not exhibit any changes in cortisol levels throughout the experiment (Fig. 4B).

**Figure 4.**
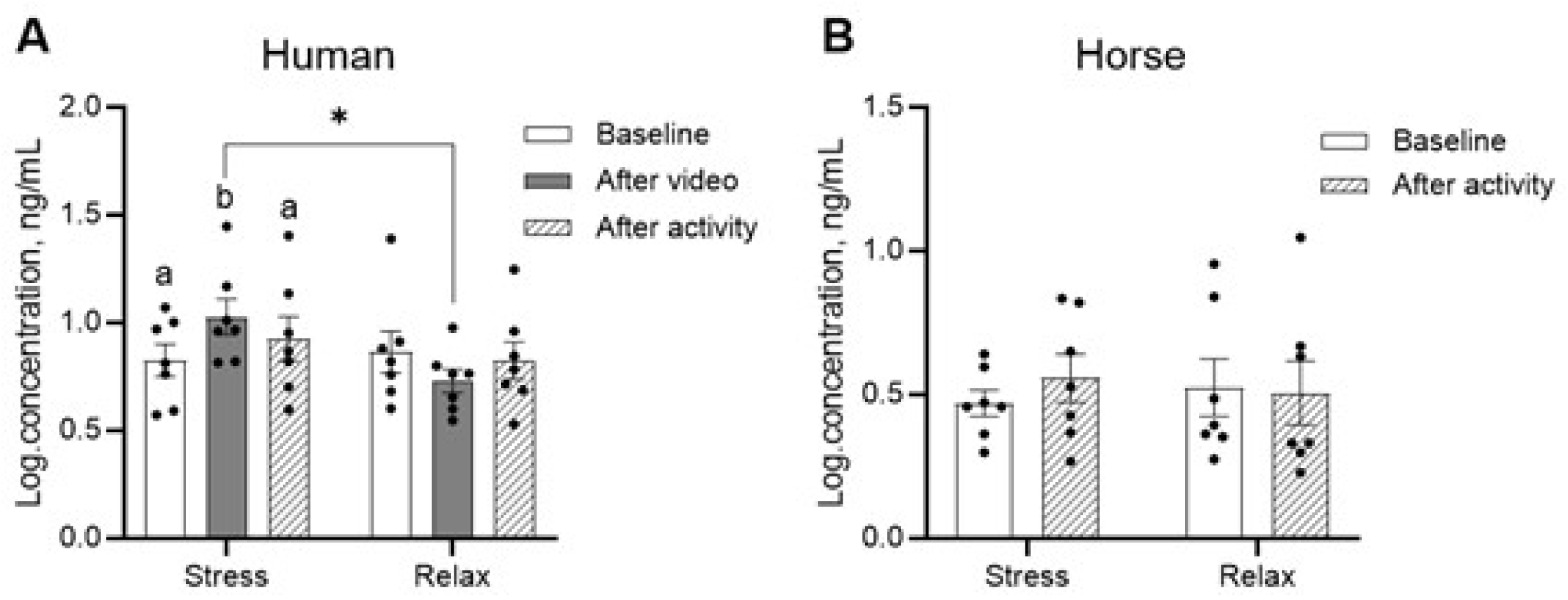
Mean concentrations of cortisol in handlers (A) and horses (B). Dots indicate individual data points. Alphabets and asterisks indicate significant differences (* < 0.05).

### 3.5. Changes in horse behaviors

Horses exhibited distinct behavioral changes depending on whether their handler had watched a stressful or relaxed video. Horses pushed the handler more frequently and showed less lip-licking behavior when the handler had watched the stressful video (Table 2). Other behaviors, such as ear pinning back, tail swishing, and bucking, did not show significant differences between conditions.

**Table 2.** The frequencies of behaviors that horses showed during interaction with handlers.

| Behaviors | Stress | Relax | *p* |
| --- | --- | --- | --- |
| Ear pinned back | 9.71±2.81 | 8.00±2.30 | 0.053 |
| Tail swishing | 16.86±7.34 | 12.14±4.57 | 0.160 |
| Push | **4.00±0.87** | **3.00±0.62** | **0.017** |
| Bucking | 0.86±0.59 | 0.14±0.14 | 0.253 |
| Resisting | 4.00±0.65 | 3.00±0.53 | 0.134 |
| Body shaking | 0.29±0.18 | 0.71±0.18 | 0.078 |
| Lip-licking | **0.29±0.18** | **1.29±0.18** | **0.003** |
| Blowing | 0.43±0.20 | 1.29±0.29 | 0.078 |

The results are expressed as the mean ± standard error of mean.

The time taken for horses to complete the tarp obstacle course and to touch the gym ball with their nose was also measured during the activities. There was no significant difference in these times between the stressed and relaxed conditions (Fig. 5), indicating that while the handler’s stress influenced certain horse behaviors, it did not affect the horses’ performance in these specific tasks.

**Figure 5.**
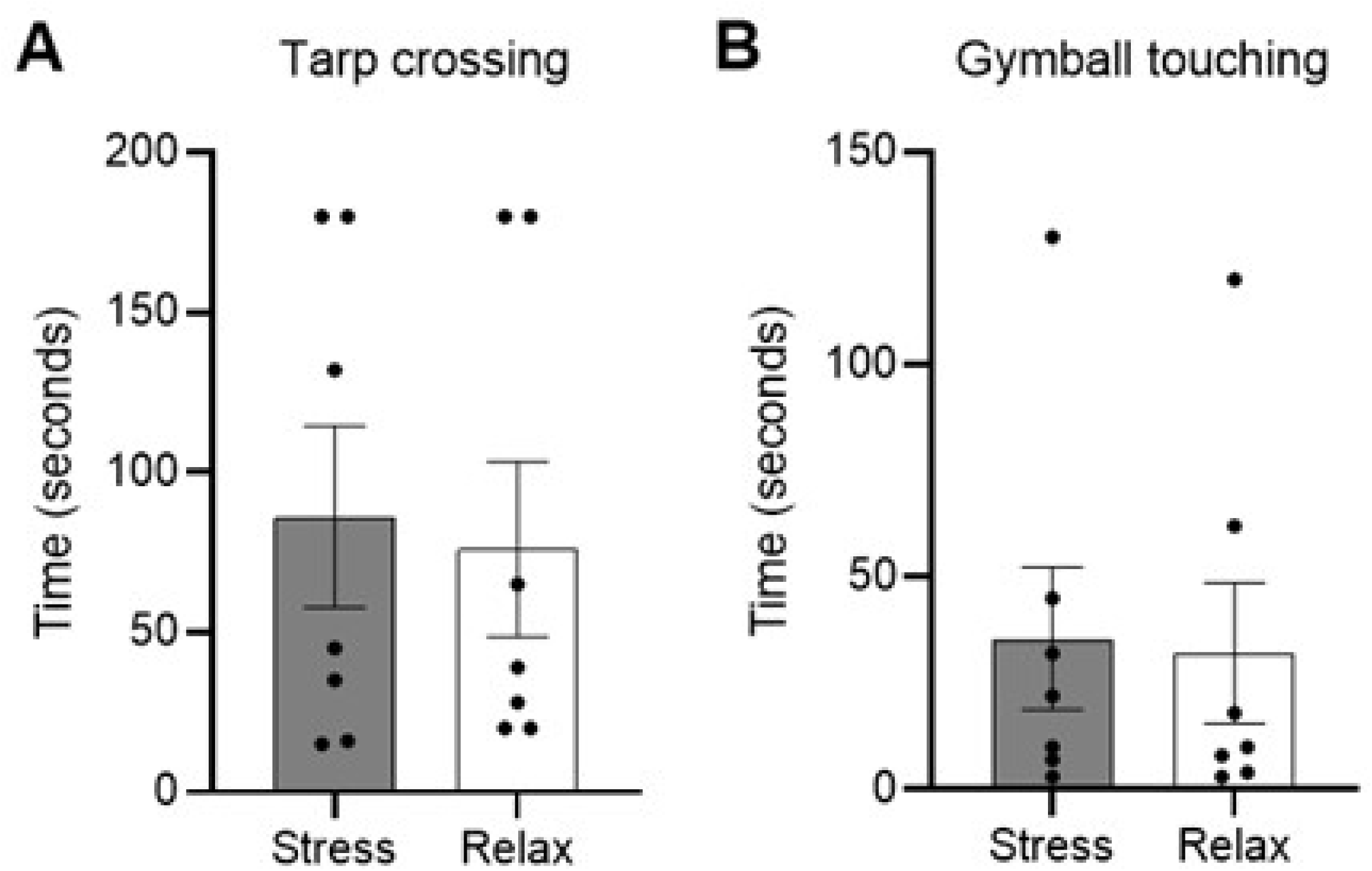
Mean seconds taken to cross over tarps (A) and touch a gym ball (B) in horses. Dots indicate individual data points.

## 4. Discussion

The findings of this study highlight the impact of stress on human brainwave activity, HRV, and cortisol levels, as well as the subsequent effects on human-animal interactions and horse behavior. Changes were most prominent in beta and high-beta brainwaves and in HRV metrics, with significant implications for understanding the bidirectional nature of stress between humans and animals.

The observed changes in brainwave activity provide insights into how stress and relaxation states affect neural processes during human-horse interactions. The significant increase in beta and high-beta waves following exposure to the stressful video is consistent with the activation of heightened arousal and cognitive alertness, which are commonly associated with stress and anxiety [28]. These findings align with previous research that associates elevated beta activity with states of stress and increased mental workload [1, 12, 15]. The decrease in beta and high-beta activity after interacting with horses suggests a potential calming effect of these interactions, supporting the hypothesis that human-horse interactions can help alleviate stress-induced neural arousal. In contrast, the lack of significant changes in alpha and theta wave activity indicates that these frequency bands may not be as sensitive to the stressors used in this study or that the stress levels induced were not sufficient to disrupt these states. Alpha waves are typically linked to relaxed and wakeful states [29], while theta waves are associated with deeper relaxation and meditative states [30]. The stability of alpha and theta activity suggests that, despite experiencing stress-induced arousal, participants did not experience a marked change in their overall state of relaxation.

The HRV results provide additional evidence of the physiological impact of stress and the mitigate effect of human-horse interactions. The increase in LF power and the LF/HF ratio after watching the stressful video indicate a shift towards sympathetic dominance, a sign of the body’s “fight or flight” response [1, 12]. This suggests that the participants experienced heightened physiological arousal, as expected during stress. Conversely, the decrease in HF power indicates a suppression of parasympathetic activity, which is usually associated with relaxation and recovery [31]. The subsequent decrease in LF power and the increase in HF power after engaging in activities with horses suggest a rebalancing of autonomic nervous system activity towards a more relaxed state. This finding supports the therapeutic potential of human-horse interactions, which may help reduce physiological stress markers and promote a calmer state [6, 16].

The HAIS results highlight the impact of human stress levels on the quality of interactions with horses. Lower HAIS scores for both humans and horses after watching a stressful video suggest that elevated human stress adversely affects these interactions. This finding indicates that horses are perceptive to their handler’s emotional states, and that human stress can lead to less favorable and cooperative behavior from the horse. This is supported by a previous study showing that horses can understand and respond to human emotions and behaviors [32]. In that study, horses comprehended and reacted to both positive and negative human emotions while watching videos and used this information to discriminate between experimenters. The sensitivity of horses to human stress highlights the importance of emotional regulation in handlers for fostering positive human-horse interactions, particularly in settings such as equine-assisted therapy or training. Conversely, higher HAIS scores following a relaxed video suggest that a calmer human state fosters more positive interactions with horses, and that horses respond more favorably to handlers who are not in a stressed state. This has practical implications for training and handling practices, suggesting that efforts to manage and reduce handler stress could enhance the quality of human-horse interactions.

The behavioral changes observed in horses provide further evidence of their sensitivity to human stress levels. The increased tendency to push handlers and the reduction in lip-licking behavior when handlers were stressed suggest that horses were more agitated or uncomfortable in these interactions. Pushing behavior indicates a desire to establish a space or assert control, while lip-licking is often associated with processing and relaxation [33, 34]. The increased pushing behavior could reflect an attempt to either distance themselves from the stressed handler or assert control over the interaction. In contrast, the reduction in lip-licking further indicates that the horses were less at ease during these stressful interactions. Interestingly, other behaviors such as ear pinning, tail swishing, and resisting did not show significant differences between conditions. The absence of significant differences in these behaviors suggests that horses may express their sensitivity to human stress in more subtle or specific ways, rather than through overt signs of distress. This implies that not all stress-related behaviors are triggered equally depending on individual’s stress coping strategies [35].

The performance of horses on tasks such as navigating an obstacle course or touching a gym ball remained consistent regardless of the handlers’ stress level. This result suggests that while handlers’ stress influenced certain interaction behaviors, it did not affect the ability of horses to perform specific tasks. This indicates that the stress in human may have a more noticeable effect on more nuanced aspects of horse behavior during direct interactions, such as subtle communication cues or responsiveness. The association between stress level and performance outcomes in horses varied depending on sports and circumstances [36–38]. This could be due to varying levels of intensity, complexity of tasks, or the specific training and conditioning the horses undergo different types of activities.

This study demonstrates the significant influence of human stress on both neural and behavioral responses during interactions with horses. Increased beta and high-beta wave activity in response to stress, along with changes in HRV, highlight the physiological effects of stress and the potential calming influence of horse interactions. The negative impact of human stress on human-horse interaction quality and horse behavior underscores the importance of managing handler’s stress to promote positive and cooperative interactions. These findings have important implications for the use of horses in therapeutic settings, equine training, and welfare practices, emphasizing the need for handlers to be aware of their emotional states and the potential effects on their equine partners.

## Acknowledgements

The authors would like to thank Dr. Heejun Jung (Korea Polytechnics, Republic of Korea), Geumhui Lee (Korea Racing Authority, Republic of Korea), Shakeel Hafiz Muhammad (Arid Agriculture University, Islamic Republic of Pakistan), Junyoung Kim, Yubin Song, Jaewoo Choi and Yujin Song (Kyungpook National University, Republic of Korea) for their support.

## Author contributions

**Conceptualization**: Yeonju Choi, Minjung Yoon

**Data curation**: Yeonju Choi

**Formal analysis**: Yeonju Choi

**Funding acquisition**: Minjung Yoon

**Investigation**: Yeonju Choi

**Methodology**: Yeonju Choi

**Project administration**: Minjung Yoon

**Resources**: Yeonju Choi, Minjung Yoon

**Software**: Yeonju Choi

**Supervision**: Carissa L. Wickens, Minjung Yoon

**Validation**: Yeonju Choi, Youngwook Jung

**Visualization**: Yeonju Choi

**Writing – original draft**: Yeonju Choi

**Writing - review & editing**: Youngwook Jung, Carissa L. Wickens, Minjung Yoon

## References

1. Arrazola A, Merkies K. Effect of human attachment style on horse behaviour and physiology during equine-assisted activities–A pilot study. Animals. 2020;10(7):1156.

2. Dashper K. Human-animal relationships in equestrian sport and leisure: Routledge; 2016.

3. Danby P, Dashper K, Finkel R. Multispecies leisure: Human-animal interactions in leisure landscapes. Taylor & Francis; 2019. p. 291–302.

4. Nakamura K, Takimoto-Inose A, Hasegawa T. Cross-modal perception of human emotion in domestic horses (Equus caballus). Scientific Reports. 2018;8(1):8660.

5. Scopa C, Contalbrigo L, Greco A, Lanatà A, Scilingo EP, Baragli P. Emotional transfer in human–horse interaction: new perspectives on equine assisted interventions. Animals. 2019;9(12):1030.

6. Kemeny ME. The psychobiology of stress. Current directions in psychological science. 2003;12(4):124–9.

7. Barker SB, Wolen AR. The benefits of human–companion animal interaction: A review. Journal of veterinary medical education. 2008;35(4):487–95.

8. Hemsworth P, Barnett J. Human-animal interactions and animal stress. The biology of animal stress: basic principles and implications for animal welfare: CABI Publishing Wallingford UK; 2000. p. 309–35.

9. Hausberger M, Roche H, Henry S, Visser EK. A review of the human–horse relationship. Applied animal behaviour science. 2008;109(1):1–24.

10. Schütz K, Rötters A, Oebel L. Individual Responses in the Domestic Horse Regarding Human Behaviour in Identical Settings. 2019.

11. Teplan M. Fundamentals of EEG measurement. Measurement science review. 2002;2(2):1–11.

12. Aris SAM, Lias S, Taib MN, editors. The relationship of alpha waves and theta waves in EEG during relaxation and IQ test. 2010 2nd International Congress on Engineering Education; 2010: IEEE.

13. Norhazman H, Zaini N, Taib MN, Jailani R, Latip MFA, editors. Alpha and beta sub-waves patterns when evoked by external stressor and entrained by binaural beats tone. 2019 IEEE 7th Conference on Systems, Process and Control (ICSPC); 2019: IEEE.

14. Choi Y, Kim M, Chun C. Measurement of occupants’ stress based on electroencephalograms (EEG) in twelve combined environments. Building and Environment. 2015;88:65–72.

15. Awang SA, Pandiyan PM, Yaacob S, Ali YM, Ramidi F, Mat F, editors. Spectral density analysis: Theta wave as mental stress indicator. Signal Processing, Image Processing and Pattern Recognition: International Conference, SIP 2011, Held as Part of the Future Generation Information Technology Conference FGIT 2011, in Conjunction with GDC 2011, Jeju Island, Korea, December 8-10, 2011 Proceedings; 2011: Springer.

16. Kaur B, Durek JJ, O’Kane BL, Tran N, Moses S, Luthra M, et al., editors. Heart rate variability (HRV): an indicator of stress. Independent Component Analyses, Compressive Sampling, Wavelets, Neural Net, Biosystems, and Nanoengineering XII; 2014: SPIE.

17. Pollard TM. Use of cortisol as a stress marker: Practical and theoretical problems. American Journal of Human Biology. 1995;7(2):265–74.

18. Hellhammer DH, Wüst S, Kudielka BM. Salivary cortisol as a biomarker in stress research. Psychoneuroendocrinology. 2009;34(2):163–71.

19. Lundberg P, Hartmann E, Roth LSV. Does training style affect the human-horse relationship? Asking the horse in a separation–reunion experiment with the owner and a stranger. Applied Animal Behaviour Science. 2020;233:105144.

20. Choi Y, Yoon M. Efficacy of androstenone in reducing stress-or fear-related responses of horses during riding. Journal of Veterinary Behavior. 2023;69:19–23.

21. Christensen JW, Uldahl M. Oral behaviour during riding is associated with oral lesions in dressage horses – A field study. Applied Animal Behaviour Science. 2024;279:106396.

22. Fureix C, Pagès M, Bon R, Lassalle J-M, Kuntz P, Gonzalez G. A preliminary study of the effects of handling type on horses’ emotional reactivity and the human–horse relationship. Behavioural Processes. 2009;82(2):202–10.

23. Warren-Smith AK, Greetham L, McGreevy PD. Behavioral and physiological responses of horses (Equus caballus) to head lowering. Journal of Veterinary Behavior. 2007;2(3):59–67.

24. Fournier AK, Berry TD, Letson E, Chanen R. The Human–Animal Interaction Scale: Development and Evaluation. Anthrozoös. 2016;29(3):455–67.

25. Abhang PA, Gawali BW, Mehrotra SC. Introduction to EEG-and speech-based emotion recognition: Academic Press; 2016.

26. Jung Y, Yoon M. The Effects of Human–Horse Interactions on Oxytocin and Cortisol Levels in Humans and Horses. Animals. 2025;15(7):905.

27. Jung Y, Yang K, Yoon M. Exploring the relationship between plasma and salivary levels of oxytocin, vasopressin, and cortisol in beagles: A preliminary study. Domestic Animal Endocrinology. 2025;92:106937.

28. Díaz H, Cid FM, Otárola J, Rojas R, Alarcón O, Cañete L. EEG Beta band frequency domain evaluation for assessing stress and anxiety in resting, eyes closed, basal conditions. Procedia Computer Science. 2019;162:974–81.

29. Sharma A, Singh M, editors. Assessing alpha activity in attention and relaxed state: An EEG analysis. 2015 1st International Conference on Next Generation Computing Technologies (NGCT); 2015: IEEE.

30. Williams JD, Gruzelier JH. Differentiation of hypnosis and relaxation by analysis of narrow band theta and alpha frequencies. International Journal of Clinical and Experimental Hypnosis. 2001;49(3):185–206.

31. Nakao M. Heart rate variability and perceived stress as measurements of relaxation response. MDPI; 2019. p. 1704.

32. Trösch M, Pellon S, Cuzol F, Parias C, Nowak R, Calandreau L, et al. Horses feel emotions when they watch positive and negative horse–human interactions in a video and transpose what they saw to real life. Animal Cognition. 2020;23(4):643–53.

33. McGreevy PD, Oddie C, Burton FL, McLean AN. The horse–human dyad: Can we align horse training and handling activities with the equid social ethogram? The Veterinary Journal. 2009;181(1):12–8.

34. McDonnell SM. The equid ethogram: a practical field guide to horse behavior: Eclipse Press; 2003.

35. Budzyńska M. Stress Reactivity and Coping in Horse Adaptation to Environment. Journal of Equine Veterinary Science. 2014;34(8):935–41.

36. Peeters M, Closson C, Beckers J-F, Vandenheede M. Rider and Horse Salivary Cortisol Levels During Competition and Impact on Performance. Journal of Equine Veterinary Science. 2013;33(3):155–60.

37. Henshall C, Randle H, Francis N, Freire R. The effect of stress and exercise on the learning performance of horses. Scientific Reports. 2022;12(1):1918.

38. Negro S, Bartolomé E, Molina A, Solé M, Gómez MD, Valera M. Stress level effects on sport performance during trotting races in Spanish Trotter Horses. Research in Veterinary Science. 2018;118:86–90.

